# Subphase-Labeled Mitotic Dataset for AI-powered Cell Division Analysis

**DOI:** 10.1101/2025.07.17.665280

**Authors:** Zsanett Zsofia Ivan, Dominik Hirling, Istvan Grexa, Jonas Ammeling, Tamas Micsik, Katalin Dobra, Levente Kuthi, Sukosd Farkas, Marc Aubreville, Vivien Miczan, Peter Horvath

## Abstract

Mitosis detection represents a critical task in the field of digital pathology, as determination of the mitotic index (MI) plays an important role in the tumor grading and prognostic assessment of patients. Manual determination of MI is a labor-intensive and time-consuming task for practitioners with rather high interobserver variability, thus, automation has become a priority. There has been substantial progress towards creating robust mitosis detection algorithms in recent years, primarily driven by the Mitosis Domain Generalization (MIDOG) challenges. In parallel, there has been growing interest in the molecular characterization of mitosis with the goal of achieving a more comprehensive understanding of its underlying mechanisms in a subphase-specific manner. Here, we introduce a new mitotic figure dataset annotated with subphase information based on the MIDOG++ dataset as well as a previously unrepresented tumor domain to enhance the diversity and applicability of the dataset. We envision a new perspective for domain generalization by improving the performance of models with subtyping mitotic cells into the 5 main stages of normal mitosis, complemented with an atypical mitotic class. We believe that our work broadens the horizon in digital pathology: subtyping information could provide useful help for mitosis detection, while also providing promising new directions in answering biological questions, such as molecular analysis of the subphases on a single cell level.

## Background & Summary

The field of digital pathology has undergone tremendous progress in recent years, for which the appearance of large vision models and the convergence of methods in the computer vision field have paved the way. Mitosis detection takes up a fairly small portion of the whole discipline, however, it has become an important area for research mainly due to its clinical relevance and the appearance of large, high quality datasets. Consequently, mitotic figure (MF) detection has become a benchmark task for domain robustness as well. The MITOS2012 challenge [1] introduced the first publicly available mitotic figure dataset. Here, domain robustness was not a priority, as the training and test sets were from the same histology slides. Subsequent challenges, such as MITOS2014 [2], AMIDA13 [3], and TUPAC16 [4], used breast cancer data and included more cases, but were limited by using only two scanners for the training and test sets. The MIDOG (MItosis DOmain Generalization) challenges were created to promote the development of domain robust detection algorithms. The MIDOG 2021 challenge [5] used a training set of 200 cases from four scanning systems and a test set of 100 cases from four scanning systems, including two previously unseen scanners. The MIDOG 2022 [5], [6] challenge included considerable diversity of tissue types and species. MIDOG++ [7] extended these datasets to include 2 mm^2^ regions of interest from 503 WSIs of seven different tumor types and labels for 11,937 mitotic figures.

Network architecture extensions with gradient reversal layers [8], [9], domain generalization capabilities of networks [10], [11], novel domain augmentation techniques [12], [13], [14] and a proper selection of unsupervised pre-training tasks [11] have all seen excessive benefit from the diverse data provided by mitosis detection research. State-of-the-art approaches exclusively use deep learning (either convolutional neural networks or vision transformers) to address the challenges of mitosis detection. Most of the published methods are framing the task as an object detection challenge, but sliding-window approaches have also been successfully applied at the cost of a higher running time [15]. The state-of-the-art MF detection method takes inspiration from the field of behavioral psychology and aims to replicate the multi-scale reasoning typical of pathologists: a “ macro-vision” step is performed which yields the semantic segmentation of mitotic candidates and a second “ micro-vision” step performs the filtering of imposter cells [11].

A significant challenge in applying mitosis detection algorithms to real-world diagnostic processes is the drop in performance when models are tested on data from different domains. Domain shifts can result from differences in species, organ type, scanner variability, or staining protocols. To address this, tissue-specific data augmentation techniques, such as stain augmentation and stain normalization, have been used to improve the generalization capabilities of mitosis detection algorithms [11], [12]. Stain augmentation can be achieved using classical methods by estimating the parameters of the staining and the scanner, or by using frequency-based approaches, such as Fourier Domain Adaptation (FDA), which swaps the low-frequency spectrum of source and target images to perform a stain adaptation between different scanners [13], [14]. Besides staining-related augmentation techniques, modifications to network architectures also proved to be an effective way to achieve domain generalization: the reference algorithm for the MIDOG2021 challenge employed a domain-adversarial training technique, which involves adding a domain classification branch to the network to encourage domain-invariant features [9].

The assessment of atypical mitotic figures (AMFs) has also gained attention as a potential prognostic marker in breast cancer. The AMi-Br [16] is one of the first datasets to include subtype information related to mitotic figures. A hierarchical anchor-free detection method has been developed to solve the mitotic vs. AMF subtyping problem [17] and the results imply that performing the classification of mitotic cells helps to improve the performance of the original MF detection task. To our knowledge, there is not yet any publicly available dataset or study that tackles the question of what happens if the mitotic cells are subtyped into the 5 different stages of normal mitosis (pro-, prometa-, meta-, ana-, and telophase) with atypical mitotic figures as a separate class.

Despite all the advancements in the research of mitosis, current methods often treat mitotic figures as a single category, overlooking potential subtypes that may hold clinical significance such as the number of atypical MFs, that have been associated with adverse prognostic factors, including reduced overall survival and poorer clinical outcomes [16], [18], [19]. In addition to this, the classification of mitosis can open up further relevant questions in research, in particular when complemented by molecular analysis [20], [21], [22]. Recent studies suggest that subtyping mitotic figures could lead to better model generalization and improved interpretability [17], yet publicly available datasets incorporating such annotations remain scarce. Moreover, while modern network architectures and augmentation strategies have pushed detection performance forward, there remains room for improvement in both object detection and classification strategies, particularly in capturing the ordinal relationships between different mitotic phases.

To address these problems, we propose a new approach that pushes existing mitosis detection workflows forward in two key areas:

- We create an annotated dataset extending MIDOG++ with detailed subtyping information. In addition to bounding boxes, we provide corresponding segmentation masks to achieve more precise localization. The necessity of segmentation is on one hand supported by the potential need for morphological analysis of the dividing cells for grading purposes. Another use-case that we envision is the isolation and molecular analysis of the mitotic cells via laser microdissection, for which contours are necessary [23]. Furthermore, we release an additional benchmarking dataset created from human lung adenocarcinoma, since it is the most common histological subtype of lung cancer with a heavy mortality burden. We publicly release the whole dataset for research purposes (Fig. 1).
- We develop a novel mitosis detection and segmentation pipeline. Our segmentation framework is built upon Mask R-CNN, a region proposal-based approach. We replace the traditional ResNet-50 backbone with ConvNeXt, a state-of-the-art feature extractor and apply a hierarchical candidate refinement step. Our pipeline also incorporates domain augmentation strategies during training.

**Fig. 1:**
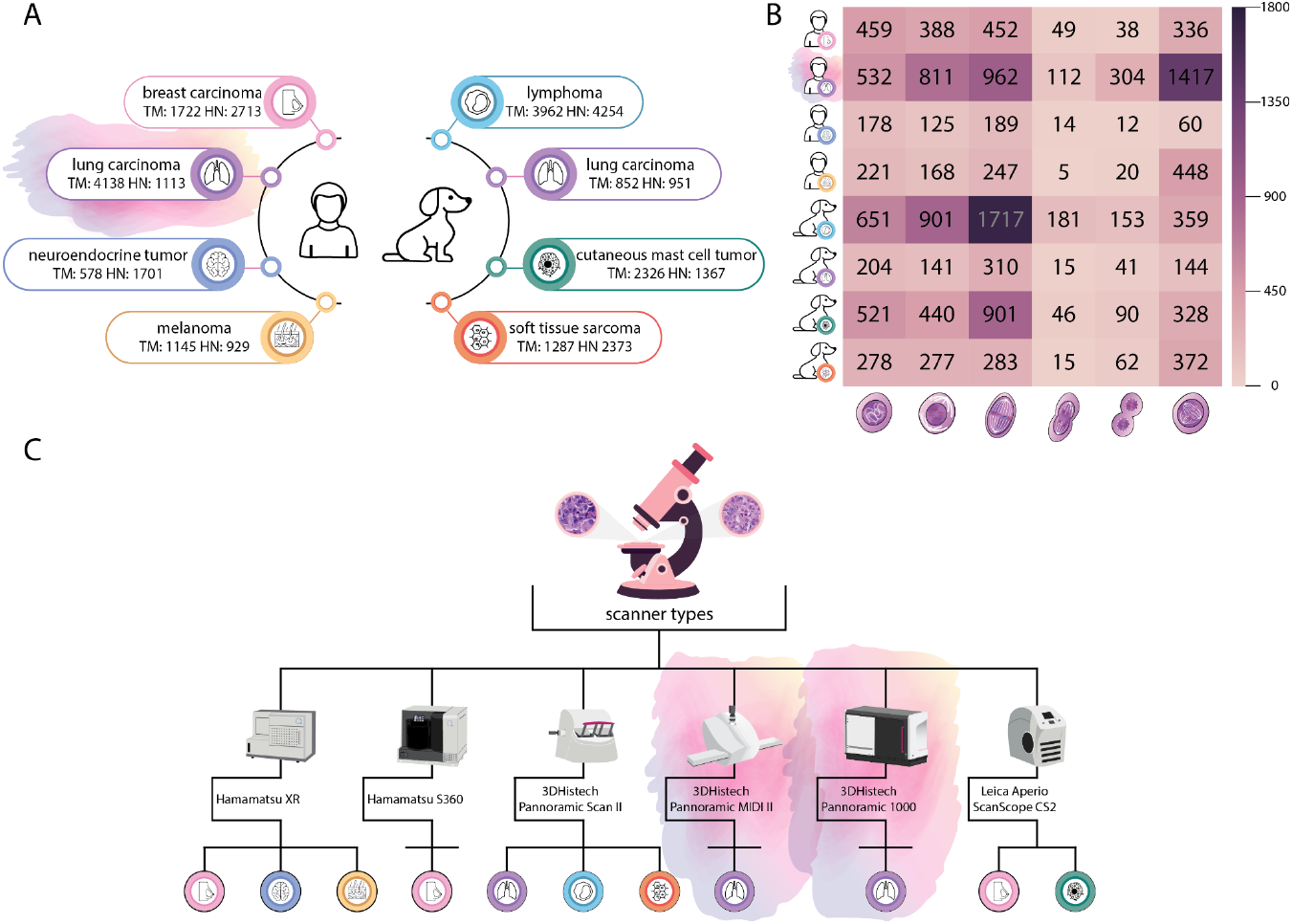
A comprehensive summary of data across species, tissues, tumor types, microscopes and mitotic figure numbers, highlighting our contribution. A) Overview of available mitotic datasets categorized by species, tumor type, and tissue of origin, complemented by the total mitotic (TM) and hard negative (HN) figure numbers B) Comparative heatmap with mitotic figure number distribution (normal prophase, prometaphase, metaphase, anaphase, telophase, and atypical mitosis, respectively) of the different tumor types C) Summary of scanner models and the corresponding tissue specimens imaged using each device

## Methods

### Specimen preparation and digitalization

For the LUNG-MITO dataset (approved under ethical license number BM/22651-1/2024 issued by the Scientific and Research Ethics Committee of the Medical Research Council -ETT TUKEB) 12 whole slides from 2 patients were selected where the tumor region was prominent. All tumor samples underwent routine diagnostic evaluation at the Department of Pathology, Faculty of General Medicine, Szent-Györgyi Albert University of Szeged. From archived formalin-fixed paraffin-embedded tissue blocks, 5 μm sections were prepared and hematoxylin-eosin staining was performed. All samples were prepared using routine pathology protocols (see Extended Data Table 1). Of the two different cases, one was scanned with 3DHISTECH PANNORAMIC MIDI II (0.24 µm/pixel) and one with 3DHISTECH PANNORAMIC 1000 (0.12 μm/pixel). In all cases the standard settings were used with 40× magnification.

### Expert labelled dataset description

#### LUNG-MITO dataset

Our mitotic figure dataset consists of human lung adenocarcinoma samples from 2 different patients and 12 slides. 1 slide from patient 1 scanned with 3DHISTECH PANNORAMIC MIDI II and 11 slides from patient 2 scanned with 3DHISTECH PANNORAMIC 1000. The images had a resolution of 0.24 and 0.12 µm/px, respectively and were tiled to 1024x1024 parts. The whole tumor area where higher mitotic activity was expected was selected for analysis as a region of interest (ROI) from each whole slide image (WSI), in order to include all potential mitotic figure candidates, not only the ones in hot spot regions, as the usual pathology protocol suggests [6], [24], [25], [26], [27]. Then, mitotic cells were selected, segmented manually and assigned into classes of the 5 major normal mitotic phases (1-Prophase, 2-Prometaphase, 3-Metaphase, 4-Anaphase, 5-Telophase) complemented by an atypical mitotic figure (AMF) class according to the previously established standards [18], [24], [28] (also see Extended Data Figure 1 and Extended Data Table 2 for our annotation strategy) using the Napari annotation software AnnotatorJ plugin [29]. To aid the deep learning models, hard negative figures were selected based on their morphological characteristics described in [18], [24], [28] (Fig. 2). Annotations are made under the supervision of expert pathologists (K.D., T.M., L.K.)

**Fig. 2:**
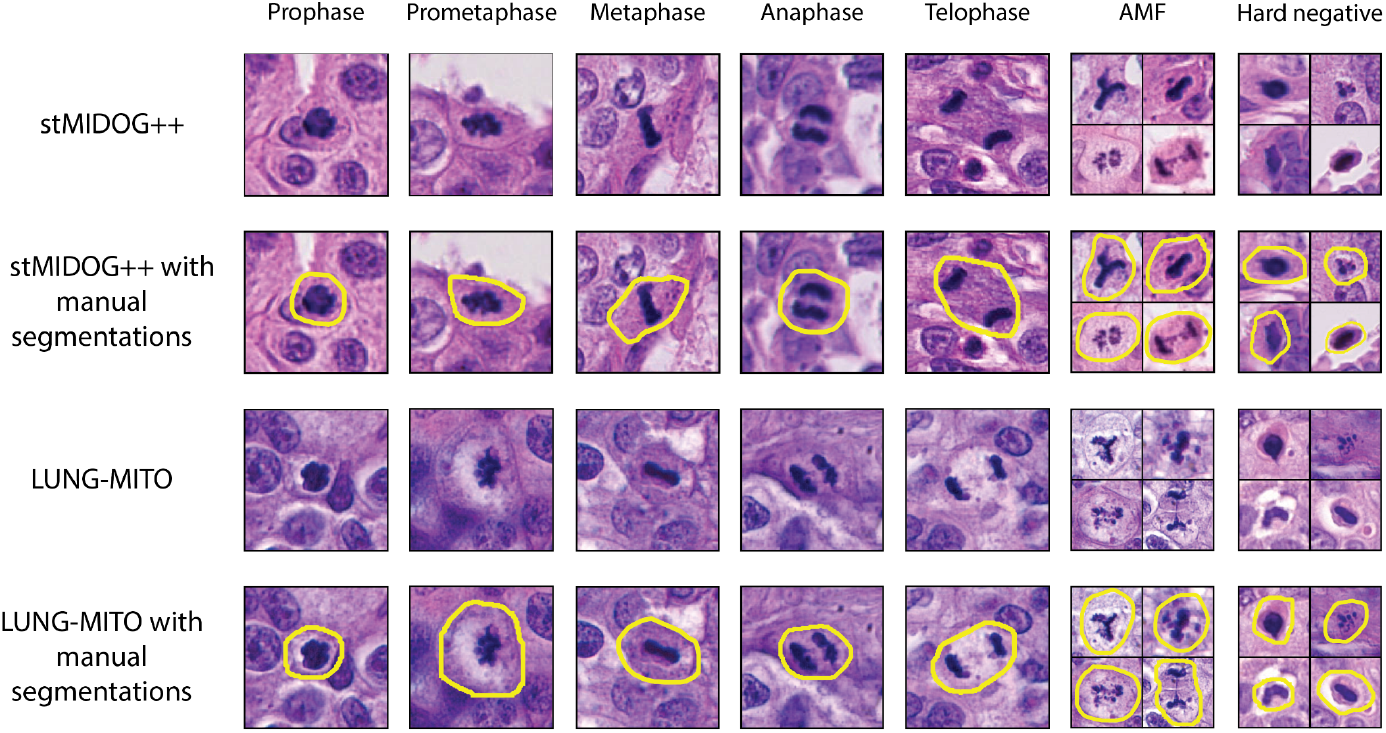
Representative images from the stMIDOG++ and the new LUNG-MITO datasets with manual segmentations. Column 1-5. Normal mitotic figures. Column 6. AMF (atypical mitotic figures) Column 7. Hard negative examples Magnification: 40× For detailed information see Extended Data Table 2

#### stMIDOG++ dataset

A large-scale annotated mitosis dataset with a total of 503 tumor cases from 10 different tumor domains (breast carcinoma, lung carcinoma, lymphoma, cutaneous mast cell tumor, neuroendocrine tumor, soft tissue sarcoma, melanoma), resolution ranging from 0.23 µm/px to 0.25 µm/px. The original dataset used 2 classes: mitotic cells and so-called hard negative instances, which are morphologically similar to mitotic cells, and a 3 expert consensus was provided, represented by bounding box coordinates. Here, we extend this labeled dataset by assigning a class from five common subphases (1-Prophase, 2-Prometaphase, 3-Metaphase, 4-Anaphase, 5-Telophase) and an atypical mitotic figure class (without further classification) to each previously annotated mitotic figure. This step was performed by a trained expert with more than 4 years of experience in labeling and identifying mitotic figures on tissue sections. Manual classification was carried out in Napari AnnotatorJ according to previously defined standards (see Extended Data Figure 1 Extended Data Table 2). In addition to subtyping, precise manual segmentations were also drawn for each of the cells (Fig. 2). This allows morphological analysis of the candidates or to proceed with single-cell isolation.

### Model description

Our mitosis detection pipeline (Fig. 3.) consists of 2 major steps: initial segmentation and candidate refinement.

**Fig. 3:**
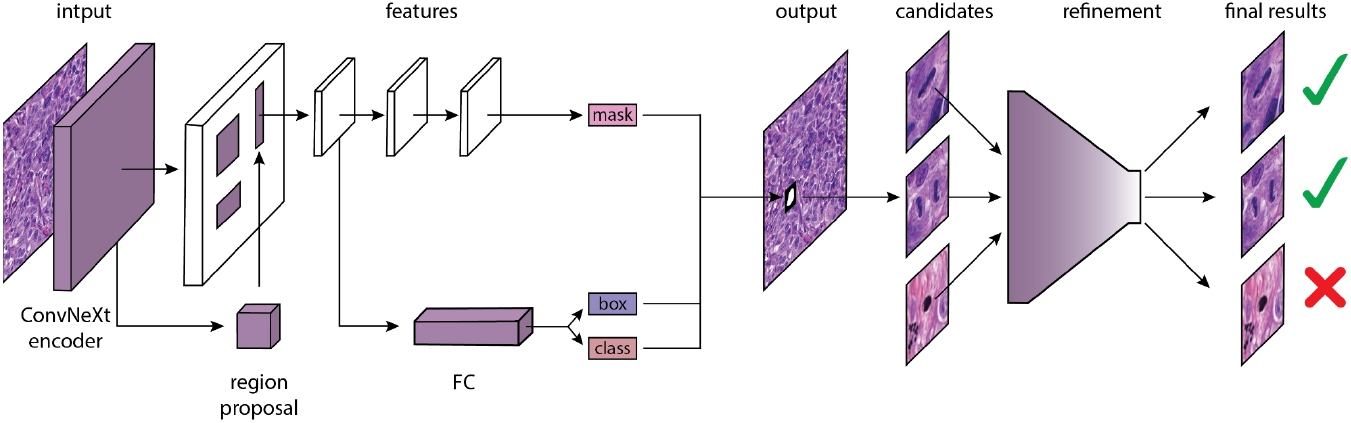
Overview of the proposed two-stage method. First, we propose mitotic candidates with a ConvNeXt encoder-based Mask R-CNN. The candidates are reclassified with EfficientNet in the second step.

#### Initial segmentation

Our segmentation network is based on Mask R-CNN [30], a popular deep learning architecture for instance segmentation. Most of the time, traditional backbones such as ResNet-50 or ResNet-101 are used [31], however, recent advancements in vision transformer-inspired architectures have demonstrated superior feature representation capabilities [32], [33]. In our approach, we replace the conventional ResNet-50 backbone with ConvNeXt, a state-of-the-art convolutional neural network that incorporates principles from vision transformers while retaining the efficiency of CNNs [34]. The output of our initial method are segmentations and subclasses of mitotic cells.

#### Candidate refinement

To further enhance detection performance, we run an additional classification step on the mitotic candidates yielded by the initial segmentation step. This step is done with EfficientNet, a state-of-the-art classification network [34], [35]. The refinement is performed in a hierarchical manner, i.e. first, a mitotic vs. imposter differentiation is performed and in the second step, the final subphase classification is done on the mitotic cells. An initial decision from Mask R-CNN is only overwritten if its confidence is lower than that of the EfficientNet.

### Evaluation methods

For evaluating the efficiency of our algorithm, the F1 score is calculated in the same manner as in the MIDOG competitions: a segmentation is counted as true positive (TP), if the centroid of the prediction is matched to a ground truth (GT) segmentation within its pixel radius *r* (for the sake of consistency, we opted for *r=25* similarly to the MIDOG evaluation metric). A ground truth object can only have one corresponding match, every other prediction will be counted as false positive (FP). Those GT segmentations that don’ t have any corresponding matches will be counted as false positive (FP).

Because of the imbalanced number of mitotic cells on the images, we aggregate our results across all images and calculate the final metric in the end:

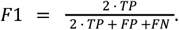

Besides a class-agnostic score, we also report class wise metrics, where TP, FP, FN samples are yielded based on detection and classification accuracy..

## Data records

The images of MIDOG++ are publicly available at the MIDOG++ GitHub Repository (https://github.com/DeepMicroscopy/MIDOGpp), while the LUNG-MITO dataset, containing 2916 PNG images, is uploaded as a zipped archive to Zenodo (https://doi.org/10.5281/zenodo.15854453).

The dataset consists of two types of data: images and the corresponding annotations in COCO format [36]. The images are 1024×1024 RGB tiles from HE-stained tumor regions from two lung adenocarcinoma patients.

The COCO annotation files include image metadata, object-level segmentation masks, bounding boxes, and category definitions. In the stMIDOG++ dataset, we preserved the original filenames from MIDOG++. In the LUNG-MITO dataset, the images are anonymized and named with numeric identifiers (e.g., 00002.png). In both COCO files, each image entry includes a unique image_id used to associate annotations with their respective images.

We provide two annotation files: *MIDOGpp_subtyping*.*json* for the stMIDOG++ dataset, containing 26283 annotations for 503 images, and *Lung_mito*.*json* for the LUNG-MITO dataset, containing 5251 annotations for 2916 images. Each object is labeled with a category ID from 1 to 7, corresponding to the following categories: 1 – prophase, 2 – prometaphase, 3 – metaphase, 4 – anaphase, 5 – telophase, 6 – non-mitotic (negative), and 7 – atypical mitotic figures.

In the LUNG-MITO dataset, the number of annotations per category is: 532 (Category 1), 811 (Category 2), 962 (Category 3), 112 (Category 4), 304 (Category 5), 1113 (Category 6), and 1417 (Category 7). In the MIDOG++ dataset, the corresponding counts are: 2522 (Category 1), 2458 (Category 2), 4112 (Category 3), 327 (Category 4), 417 (Category 5), 14347 (Category 6), and 2100 (Category 7).

In the COCO annotation files, segmentation masks are encoded as lists of polygon points in the format [x0, y0, x1, y1, …, xn, yn], and bounding boxes are specified using the [x_min, y_min, width, height] format.

The source code is available at: (https://github.com/biomag-lab/Mitosis-detection).

## Technical validation

The datasets were partitioned as follows: a subset of stMIDOG++ was used exclusively for training, while the LUNG-MITO dataset was reserved for evaluation to assess the domain robustness of the algorithm. 70% of the stMIDOG++ dataset’ s images were allocated for training, 10% for validation (used for confidence threshold optimization and hyperparameter tuning), and 20% for testing. To enable fair evaluation, a 3-fold cross-validation was performed using the training and validation sets, with the test set kept fixed. All performance metrics were computed as the average across the three folds. To ensure an unbiased evaluation, domain leakage between training, validation, and test sets was avoided by preventing overlap of crops at the slide level. Although some tissue types may appear in both training and testing sets, the inclusion of the LUNG-MITO dataset ensures that domain robustness is rigorously assessed.

To benchmark the performance of the algorithm, comparisons were made against a classical Mask R-CNN architecture with a ResNet-50 backbone. Additionally, the contribution of the refinement step was assessed by evaluating the ConvNeXt-based method both with and without the refinement component.

All three algorithms were trained with the same hyperparameters and on the same folds. All three algorithms were based on the Mask R-CNN implementation of the MMDetection library [37].

Our results show that incorporating a ConvNeXt backbone can significantly enhance the performance of Mask R-CNN both on the stMIDOG++ and the LUNG-MITO datasets. As for the refinement part, we can see that a significant part of the additional performance is yielded by this post-processing step (Table 1).

**Table 1:** Comparison of detection performance across models. Metrics are reported as average values over three cross-validation folds.

| stMIDOG++ test set |  |  |  |  |  |  |
| --- | --- | --- | --- | --- | --- | --- |
| Model | F1 | Precision | Recall | cw-F1 | cw-Precision | cw-Recall |
| Mask R-CNN (ResNet-50) | 0,683 | 0,641 | 0,731 | 0,538 | 0,541 | 0,536 |
| Ours | 0,767 | 0,768 | 0,766 | 0,617 | 0,624 | 0,610 |
| Ours (refinement) | 0,794 | 0,774 | 0,818 | 0,630 | 0,612 | 0,650 |

| LUNG-MITO |  |  |  |  |  |  |
| --- | --- | --- | --- | --- | --- | --- |
| Model | F1 | Precision | Recall | cw-F1 | cw-Precision | cw-Recall |
| Mask R-CNN (ResNet-50) | 0,607 | 0,593 | 0,621 | 0,303 | 0,590 | 0,205 |
| Ours | 0,739 | 0,711 | 0,769 | 0,475 | 0,630 | 0,380 |
| Ours (refinement) | 0,761 | 0,748 | 0,779 | 0,483 | 0,603 | 0,404 |

## Conclusion

We present a new, publicly available dataset for mitosis detection that incorporates subphase annotations and segmentation masks, enabling both precise cell classification and potential downstream molecular analysis. By extending the stMIDOG++ dataset and introducing the LUNG-MITO benchmark set, we address gaps in domain diversity and phase-specific labeling. Our proposed detection pipeline, combining a ConvNeXt-based Mask R-CNN with hierarchical refinement via EfficientNet, achieves substantial performance gains over traditional approaches. These results confirm that mitotic subtyping not only supports improved model generalization but also opens the door for deeper biological insights. Our work establishes a new benchmark for phase-resolved mitotic figure analysis and lays a foundation for future integration of morphological and molecular cell cycle profiling.

## Code availability

Source code for model configuration, training, prediction and evaluation is available at https://github.com/biomag-lab/Mitosis-detection.

## Acknowledgments

We acknowledge support from the TKP2021-EGA09, TKCS-2024/73, Horizon-BIALYMPH, Horizon-SYMMETRY, Horizon-SWEEPICS, H2020-Fair-CHARM, HAS-NAP3, from OTKA-SNN no. 139455/ARRS and OTKA-Excellence 2025, OTKA PD 147127, and Finnish Cancer Society.

This research was supported by the Ministry for Innovation and Technology of Hungary from the National Research, Development and Innovation Fund, financed under the Cooperative Doctoral Programme funding scheme (Project no. [EKÖP-KDP-24-SZTE-11]) and the University Research Fellowship Programme funding scheme (Project no. [EKÖP-24-4-SZTE-642]).

## Author information

These authors contributed equally: Zsanett Zsófia Iván, Dominik Hirling

### Contributions

Conceptualization: Zs.Zs.I., D.H., M.A., V.M., P.H.

Data curation: Zs.Zs.I.; D.H., I.G., K.D.; T.M., L.K.

Formal analysis: D.H., I.G., J.A.

Funding acquisition: Zs.Zs.I., D.H., V.M., P.H.

Investigation: Zs.Zs.I., D.H., I.G., J.A.

Methodology: D.H., I.G.

Project administration: V.M., P.H.

Resources: Zs.Zs.I., F.S., V.M.

Software: D.H., I.G., J.A.

Supervision: V.M.; M.A., P.H.

Validation: Zs.Zs.I., D.H.

Visualization: Zs.Zs.I, V.M.

Writing - original draft: Zs.Zs.I., D.H.

Writing - review and editing: Zs.Zs.I., D.H., I.G., J.A., T.M., K.D., L.K., F.S., M.A., V.M., P.H.

## Extended data

**Extended Data Figure 1.**
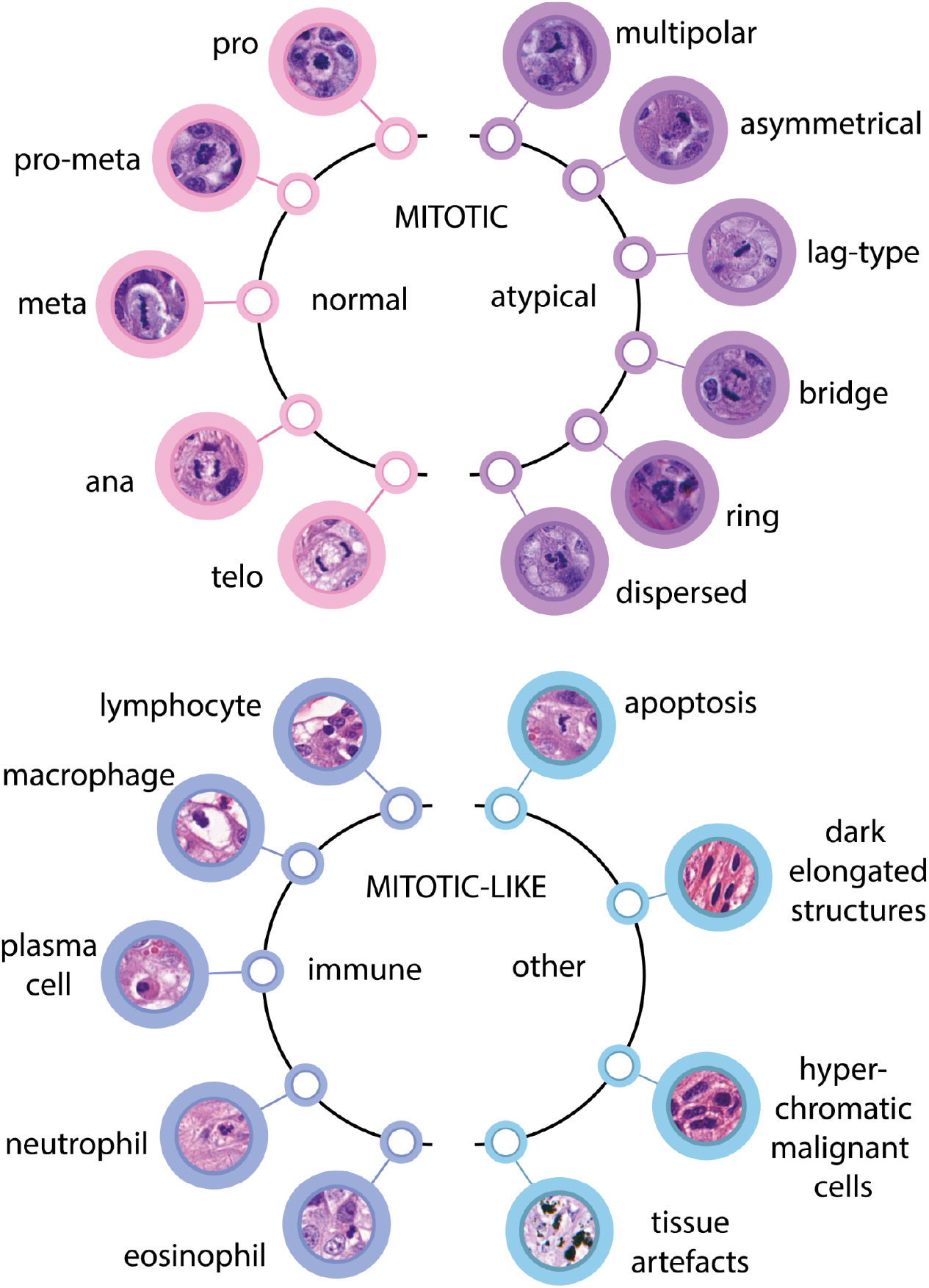
Annotation strategy with example mitotic and mitotic-like figures.

**Extended Data Table 1.** Detailed sample processing protocol.

| **Sample preparation step** | **Reagent** | **Time** | **Note** |
| --- | --- | --- | --- |
| **Tissue fixation** | 10% Formaldehyde | Overnight | Room temperature |
| **Paraffination** | 70% v/v Ethanol in water | 2x30 minutes |  |
|  | 90% v/v Ethanol in water | 30 minutes |  |
|  | 96% v/v Ethanol in water | 30 minutes |  |
|  | 2:1 propan-2-ol and 96% Ethanol in water | 3x1.5 hours |  |
|  | 1:1 propan-2-ol+Xylene | 30 minutes |  |
|  | Xylene | 2x1 hour |  |
|  | Paraffin | 4x1 hour | The sample stays in the last paraffin till the morning |
| **Tissue embedding** | Paraffin | 30 minutes | Paraffin block generation |
| **Specimen slicing with microtome** | Water bath 46°C | 10 minutes | Thickness: 5µm |
| **H&E staining** | Xylene | 2 minutes |  |
|  | Xylene | 2 minutes |  |
|  | Absolute Ethanol | 1 minute |  |
|  | 70% Ethanol in water | 1 minute |  |
|  | 50% Ethanol in water | 1 minute |  |
|  | Distilled water | 1 minute |  |
|  | Hematoxylin | 1 minute |  |
|  | Tap water | 2,5 minutes |  |
|  | Tap water | 2,5 minutes |  |
|  | Eosin Y | 30 seconds |  |
|  | 70% Ethanol in water | 1 minute |  |
|  | Absolute Ethanol | 1 minute |  |
|  | Absolute Ethanol | 1 minute |  |
|  | Xylene | 2 minutes |  |
|  | Xylene | 2 minutes |  |
| **Coverslipping with mounting medium** | Leica Surgipath Micromount | 10 minutes | 50 µl/sample<br>Size of coverslip: 22x40 mm<br>Let it cure for 20 minutes before imaging |

**Extended Data Table 2.**
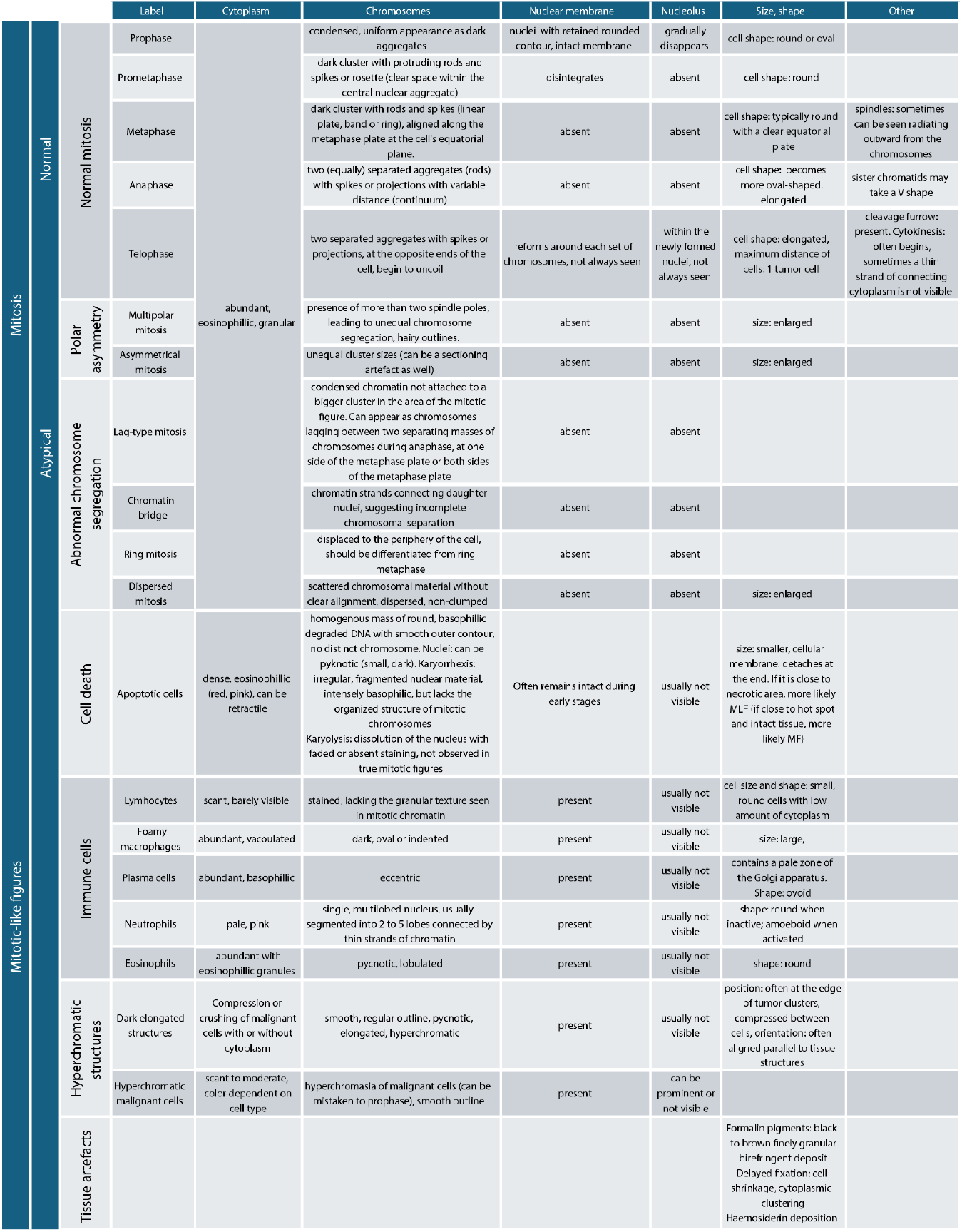
Morphological characteristics of mitotic cells and mitotic-like figures.

